# *In Situ* Visualization of the pKM101-Encoded Type IV Secretion System Reveals a Highly Symmetric ATPase Energy Center at the Channel Entrance

**DOI:** 10.1101/2021.06.01.446691

**Authors:** Pratick Khara, Peter J. Christie, Bo Hu

**Author notes:** Address correspondence to: Bo Hu, or Peter J Christie.

## Abstract

Bacterial conjugation systems are members of the type IV secretion system (T4SS) superfamily. T4SSs can be classified as ‘minimized’ or ‘expanded’ based on whether assembly requires only a core set of signature subunits or additional system-specific components. The prototypical ‘minimized’ systems mediating *Agrobacterium tumefaciens* T-DNA transfer and conjugative transfer of plasmids pKM101 and R388 are built from 12 subunits generically named VirB1-VirB11 and VirD4. In this study, we visualized the pKM101-encoded T4SS in the native context of the bacterial cell envelope by *in situ* cryoelectron tomography (CryoET). The T4SS_pKM101_ is composed of an outer membrane core complex (OMCC) connected by a thin stalk to an inner membrane complex (IMC). The OMCC exhibits 14-fold symmetry and resembles that of the T4SS_R388_, a large substructure of which was previously purified and analyzed by negative-stain electron microscopy (nsEM). The IMC of the *in situ* T4SS_pKM101_ machine is highly symmetrical and exhibits 6-fold symmetry, dominated by a hexameric collar in the periplasm and a cytoplasmic complex composed of a hexamer of dimers of the VirB4-like TraB ATPase. The IMC closely resembles equivalent regions of three ‘expanded’ T4SSs previously visualized by *in situ* CryoET, but strikingly differs from the IMC of the purified T4SS_R388_ whose cytoplasmic complex instead presents as two side-by-side VirB4 hexamers. Together, our findings support a unified architectural model for all T4SSs assembled *in vivo* regardless of their classification as ‘minimized’ or ‘expanded’: the signature VirB4-like ATPases invariably are arranged as central hexamers of dimers at the entrances to the T4SS channels.

**Significance:** Bacterial type IV secretion systems (T4SSs) play central roles in antibiotic resistance spread and virulence. By cryoelectron tomography (CryoET), we solved the structure of the plasmid pKM101-encoded T4SS in the native context of the bacterial cell envelope. The inner membrane complex (IMC) of the *in situ* T4SS differs remarkably from that of a closely-related T4SS analyzed *in vitro* by single particle electron microscopy. Our findings underscore the importance of comparative *in vitro* and *in vivo* analyses of the T4SS nanomachines, and firmly support a unified model in which the signature VirB4 ATPases of the T4SS superfamily function as a central hexamer of dimers to regulate substrate entry into and passage through T4SS channels.

## Introduction

Many species of bacteria deploy type IV secretion systems (T4SSs) to deliver DNA or protein substrates to target cells (1–3). T4SSs designated as ‘minimized’ systems are assembled from a core set of signature subunits, while others termed ‘expanded’ are compositionally and structurally more complex, possibly reflecting adaptations arising over evolutionary time for specialized functions (3). In Gram-negative species, ‘minimized’ systems are assembled from ~12 subunits generically named VirB1 - VirB11 and VirD4 based on the paradigmatic *Agrobacterium tumefaciens* VirB/VirD4 T4SS (3). Three subunits (VirB7, VirB9, C-terminus of VirB10) assemble as an outer membrane core complex (OMCC) that spans the distal region of the periplasm and outer membrane (OM) (4). Four integral membrane components (VirB3, VirB6, VirB8, N-terminus of VirB10) and two or three ATPases (VirB4, VirD4, +/- VirB11) with catalytic domains located in the cytoplasm, together comprise the inner membrane complex (IMC) (5, 6). Some T4SSs elaborate an extracellular organelle termed the conjugative pilus from homologs of the VirB2 pilin and VirB5 pilus-tip subunit (3). ‘Expanded’ systems are composed of homologs or orthologs of most or all of the VirB/VirD4 subunits plus as many as 20 components that are system-specific (3).

To better understand the mechanism of action of T4SSs and the structural bases underlying functional diversity, there is growing interest in solving the structures of intact machines or machine subassemblies. OMCCs are generally stable and amenable to purification, and structures are now available for OMCCs from several ‘minimized’ and two ‘expanded’ systems at resolutions approaching ~3 Å (4, 6–12). Structural analyses of inner membrane (IM) portions of T4SSs have been considerably more challenging likely due to dissociation or distortion of these regions of T4SSs during purification. Presently, one structure exists for a ‘minimized’ system encoded by the conjugative plasmid R388. Designated the VirB_3-10_ complex, this structure was obtained by overproduction of the VirB3 - VirB10 homologs, affinity purification of the detergent-solubilized complex, and analysis by negative-stain electron microscopy (nsEM) (6). The VirB_3-10_ complex consists of the OMCC and IMC connected by a thin, flexible stalk. The IMC is composed of a highly asymmetric IM platform connected to two side-by-side hexamers of the VirB4 ATPase extending into the cytoplasm. In an updated structure, two dimers of the VirD4 ATPase were shown to integrate between the VirB4 barrels (13).

IMCs of ‘expanded’ T4SSs have not yet been analyzed by single-particle EM. However, recent advances in *in situ* cryoelectron microscopy (CryoET) have enabled visualization of the *L. pneumophila* Dot/Icm, *H. pylori* Cag, and F plasmid-encoded Tra T4SSs (hereafter designated T4SS_Dot/Icm_, etc.) in the native context of the cell envelope (14–21). Remarkably, in contrast to the IMC of the VirB_3-10_ structure, the IMCs of all three of the ‘expanded’ systems clearly exhibit 6-fold symmetry and the VirB4 ATPases instead assemble as a central hexamer of dimers at the channel entrance (16–18).

In the present study, we solved the *in situ* structure of the pKM101-encoded T4SS (hereafter T4SS_pKM101_), which is phylogenetically (**Fig. S1A**) and functionally closely related to the R388-encoded T4SS (T4SS_R388_) to the extent that the two ‘minimized’ systems translocate each other’s plasmids and some machine subunits are swappable (8). We report that the IMC of the *in situ* T4SS_pKM101_ adopts the 6-fold system observed for the equivalent regions of the ‘expanded’ T4SSs. Most strikingly, the VirB4 homolog is arranged as a central hexamer of dimers, not the side-by-side hexameric barrels visualized for this ATPase in the purified VirB_3-10_ complex. Our findings support a unified structural model for the IMCs of T4SSs regardless of their classification as ‘minimized’ or ‘expanded’: in their native membrane environments these subassemblies are highly symmetric and the signature VirB4 ATPases invariably adopt a hexamer of dimer configuration at the T4SS channel entrances.

## Results and Discussion

### *In situ* detection of the pKM101 nanomachine

To visualize T4SS_pKM101_ nanomachines, we deployed an *E. coli mreB minC* mutant carrying pKM101 to generate small (<300 nm in diameter) minicells (17). Minicells are ideal for *in situ* CryoET because of their small size and full metabolic capacity, including the capacity to function as donors of the pKM101 plasmid to recipient cells **(Fig. S1B)** (22). We used a high-throughput CryoET pipeline to visualize thousands of *E. coli* minicells (see **Fig. S2** for workflow). The pKM101 nanomachines were smaller and more difficult to detect than the F plasmid-encoded T4SS or other ‘expanded’ systems, but we were able to detect ~1 or 2 pKM101-encoded structures for every 2 or 3 minicells examined (**Figs. 1Ai, iii, Movie S1**). Importantly, minicell preparations from the parental strain UU2834 alone lacked these surface structures, confirming that the presence of pKM101 is required for their elaboration.

**Fig. 1.**
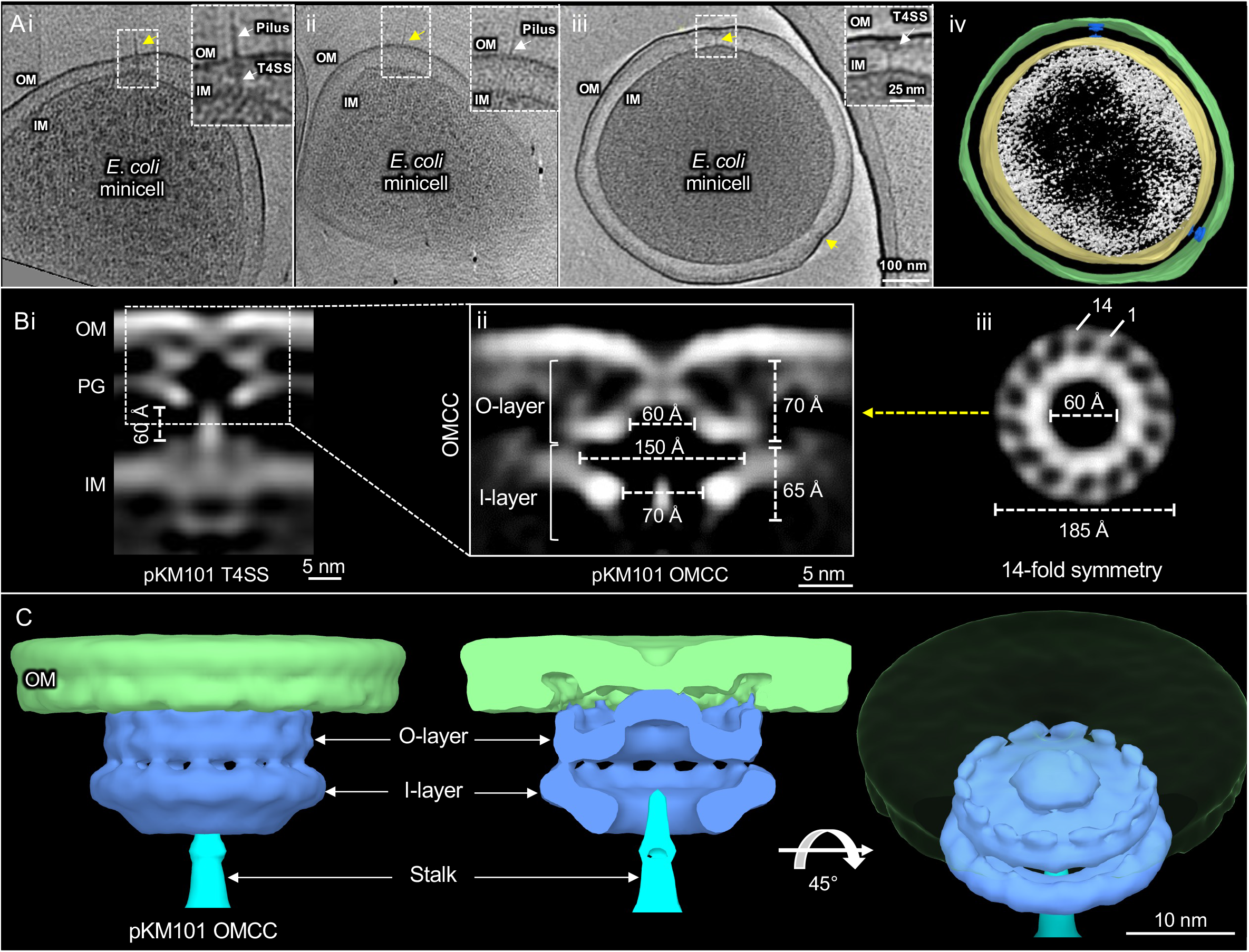
*E. coli* minicells carrying pKM101 encoded type IV secretion system (T4SS_pKM101_) and *in situ* structure of the outer membrane core complex (OMCC) of T4SS_pKM101_ revealed by CryoET and subtomogram averaging. **(A i, ii, iii)** Tomographic slices from representative *E. coli* minicells showing T4SSs embedded between the outer membrane (OM) and inner membrane (IM). pKM101 pili were associated with a few visualized T4SSs (Ai), although pilus associated OM structures without any periplasmic densities. The T4SS and novel structures are marked with yellow arrow. The boxed regions was magnified to show T4SS with and without associated pilus and also pilus associated OM structures. (A iv) A 3D surface view of the *E. coli* minicell in A iii showing T4SSs. (B i) A central slice of the averaged structure of the T4SS in the cell envelope. (B ii) After refinement, details of the OMCC are visible. The widths and heights of O-layer and I-layer chambers are shown. (B iii) A cross-section view of the region in B ii marked by a yellow arrow shows 14-fold symmetry of the OMCC. (C) 3D surface renderings of the OMCC is shown in different views.

The pKM101-encoded structures characteristically consisted of periplasmic cone-shaped complexes alone or in association with thin ‘stalk-like structures that extended to the IM (**Fig. S2**). Although the former represented about half of the initially ‘picked’ structures, they were discarded as possible assembly intermediates or dead-end complexes (**Fig. S2**). The pKM101 transfer system also elaborates conjugative pili, and accordingly we detected some pKM101 pili on the surfaces of minicells. Interestingly, these pili were bound either to the presumptive T4SS_pKM101_ machines or to sites on the OM that lacked any underlying basal structures (**Figs. 1Ai, ii, S3**). The pKM101 pili were rarely detected, however, likely because pKM101 and related IncN plasmids elaborate rigid, brittle pili that are readily detached or sloughed from cells (23). Consequently, here we focused on solving the structures of T4SS_pKM101_ machines provisionally composed of the previously-described OMCC and IMC subassemblies (6).

### Visualization of the *in situ* OMCC_pKM101_

From 287 nanomachine subtomograms extracted from 560 tomographic reconstructions, we generated *in situ* structure of the OMCC_pKM101_ at a resolution of ~37 Å (**Figs. 1B, S2**). Three-dimensional classifications revealed 14-fold symmetrical features of the OMCC which were resolved further by imposing a 14-fold symmetry during refinement **(Fig. S2)**. In the refined structure, the OMCC is clearly seen attached to the OM where it causes an invagination of the outer leaflet (**Figs. 1Bi, ii).** The upper region, designated as the O-layer in accordance with a nomenclature used to describe the *in vitro* OMCC_pKM101_ structure (4), is 185 Å wide and 70 Å in height. In side-view, the complex forms at least two contacts with the OM, the first mediated by a central cap and the second by the periphery of the OMCC (**Fig. 1Bii**). In the middle of the central cap and extending across the OM is a region of lower density that might correspond to the OM-spanning portion of the translocation channel. In top-down view, the periphery of the OMCC clearly consists of 14 knobs arranged in a ring of ~185 Å (**Fig. 1Biii**). The knobs are connected via spokes to a central contiguous ring that conforms to the base of the cap (**Fig. 1Biii**). In the 3D renderings, it is evident that the 14 peripheral knobs interact with the OM (**Fig. 1C**). Notably, besides invagination of the OM at the cap junction, the region of the OM delimited by the cap and OMCC peripheral contacts lacks an inner leaflet, suggestive of extensive remodeling of the OM during machine biogenesis **(Figs. 1Bii, iii).**

The chamber within the O-layer widens to ~150 Å at the junction with the lower region of the OMCC, which we termed the I-layer again in accordance with a previous nomenclature (4). The I-layer has a height of 65 Å and is slightly wider than the O-layer, although the outer boundary of the O-layer is blurred because of density contributed by the peptidoglycan (PG) layer. The I-layer narrows at its base, and at this position the central chamber has a diameter of ~70 Å. A ‘stalk-like’ density embeds into the central cavity and projects through the periplasm to the IM (**Fig. 2B, D**). Combined, the O- and I-layers yield a total height of 135 Å for the *in situ* OMCC structure (**Fig. 2B, D**).

**Fig. 2.**
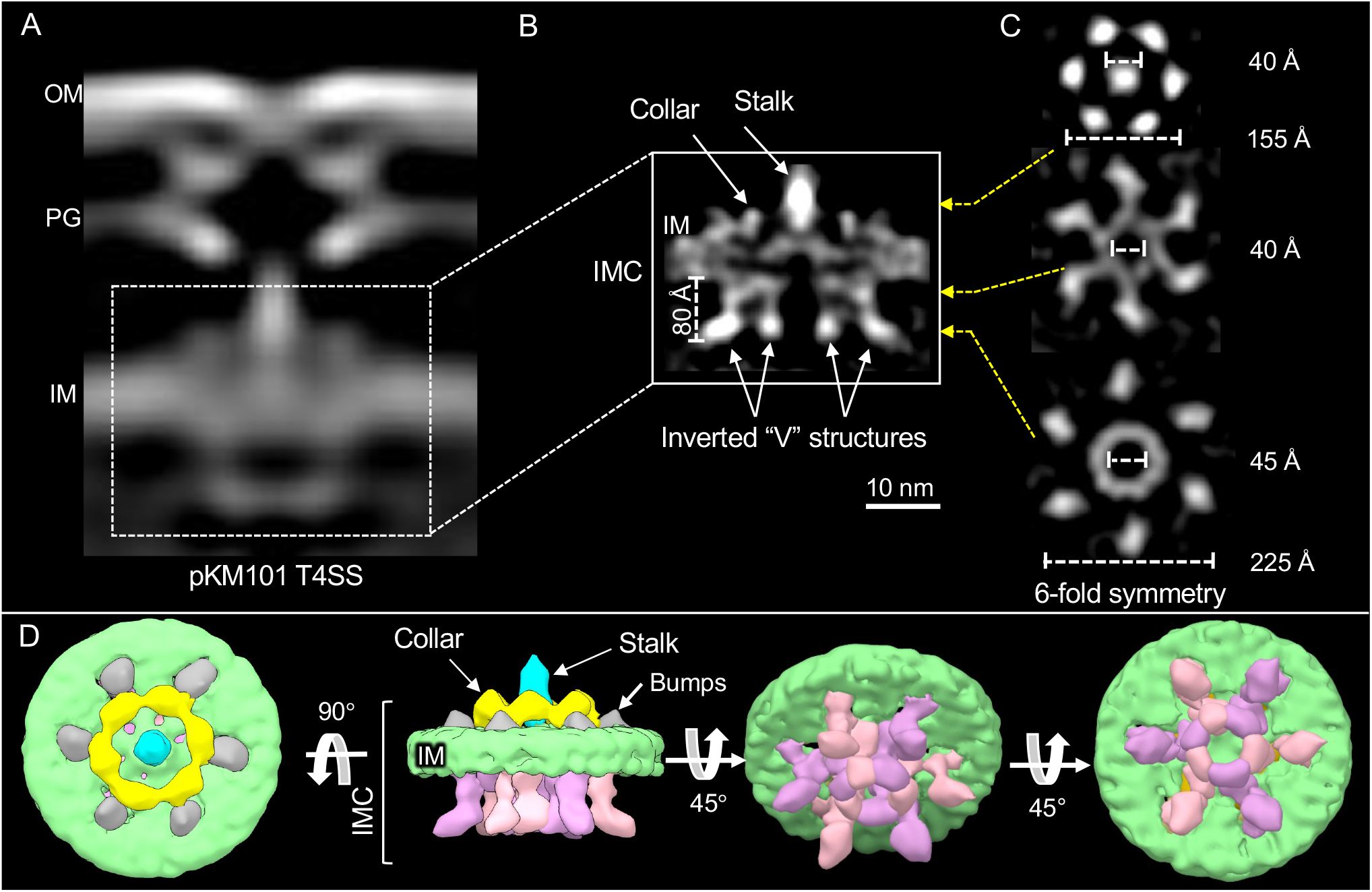
*In situ* structure of the inner membrane complex (IMC) of pKM101 encoded type IV secretion system (T4SS_pKM101_) revealed by CryoET and subtomogram averaging. (A) A central slice of the averaged structure of the T4SS in the cell envelope. (B) After refinement, details of the IMC are visible. The stalk, collar and inverted “V” structures along with the height of its arms are shown. (C) Cross-section views of the regions in B marked by yellow arrows show 6-fold symmetry of the IMC. The collar exist as a hexameric ring like structure around the central stalk. (D) 3D surface renderings of the IMC is shown in different views.

Although the resolution of the visualized OMCC (~37 Å) is lower than that achieved by purification and single particle analyses of the corresponding complexes (<20 Å) (4, 24), the *in situ* and *in vitro* structures are highly similar (**Fig. S4**). Specifically, they exhibit 14-fold symmetry and have cross-section dimensions of 185 Å, and they are composed of distinct O- and I-layers that house large central chambers (4, 7). One difference of interest is that the *in vitro* structures are more elongated (~185Å in height) than the *in situ* structure (135 Å) (**Fig. S4B, C**). An X-ray structure generated for the O-layer of the OMCC (7) fits well into the O-layer of the *in situ* structure (**Fig. S4D**), suggesting that this region of the OMCC purifies well without major distortions. Similarly, the upper portion of the I-layer, which is composed of large, N-terminal globular domains of TraO_B9_, also has been shown to be intrinsically stable (24). By contrast, the lower portion of the I-layer is built from α-helical linker domains of TraF_B10_, which connect the OMCC to the IM (24). Although these linker domains were modeled in the *in vitro* structures (4, 24), we suspect they are too flexible for detection *in vivo* or they fold inward to generate part of the central stalk. Accordingly, the OMCC visualized *in situ* would consist of the O-layer and only the upper portion of the I-layer to yield an overall height of 135 Å.

The pKM101-encoded OMCC visualized *in situ* also fitted well on the equivalent subassembly of the VirB_3-10_ complex obtained from the T4SS_R388_ machine (**Fig. S4Ei - iii**), in agreement with earlier findings that the T4SS_pKM101_ retains function when its OMCC is swapped with that from T4SS_R388_ (8). Also of importance, the thin, central stalk visualized in the VirB_3-10_ complex is also present in the *in situ* T4SS_pKM101_ structure (**Fig. S4Ei-iii**). This confirms that the central stalk is a feature of ‘minimized’ systems assembled *in vivo*, and not a structural artefact resulting from detergent solubilization and complex purification. As shown with the VirB_3-10_ complex, the central stalk of the *in situ* T4SS_pKM101_ terminates in the I-layer and forms the only detectable density linking the OMCC with the IMC. Although, this gap is ~33Å in the VirB_3-10_ structure, we observed the distance between the OMCC and IMC of the *in situ* T4SS_pKM101_ machines to be more variable and in the range of ~50 - 70 Å. Perhaps most importantly, the central stalk lacks a detectable channel (**Figs. 1, S4Eiii**). This distinguishes the ‘minimized’ pKM101 and R388 systems from ‘expanded’ systems such as the F plasmid Tra and *L. pneumophila* Dot/Icm T4SSs, whose structures were recently solved by *in situ* CryoET (16, 17). In these latter systems, the OMCCs are invariably connected to their respective IMCs by large cylinders with clearly detectable central channels (**Fig.** S4Fi-iii).

### Visualization of the *in situ* IMC_pKM101_

Next, we refined the structure of the IMC using class averages in which machines with clearly discernible OMCC and IMC densities were detected (**Figs. 2A, B, S2**). Notable features of the IMC include a distinct collar surrounding the central stalk, which in top-down view presents as six knobs arranged in a ring of 155 Å. The collar was also flanked by six protrusions or ‘bumps’ extending from the IM into the periplasm (**Fig. 2B, D**). At the cytoplasmic face of the IM, the IMC was dominated by two side-by-side inverted ‘V’ structures with apices embedded into the IM and ‘arms’ projecting ~80 Å into the cytoplasm (**Fig. 2B**). In end-on view, six V structures clearly form two concentric circles, the outer arms of the V’s configured as a knobbed ring of ~225 Å in diameter and the inner arms joining together as a central hexameric ring with an outer diameter of ~60 Å and a lumen of ~45 Å. As with the periplasmic collar, the 6-fold symmetry of the concentric rings was readily visible among the class average images without symmetry imposed (**Fig. S2**). The structure was resolved further by imposing a 6-fold symmetry during refinement (**Figs. 2B–D, S2**).

The structure of the IMC_pKM101_ bears striking similarities to IMCs associated with the ‘expanded’ F plasmid-encoded Tra and *L. pneumophila* Dot/Icm systems whose structures also were solved by *in situ* CryoET **(Fig. 3A–C)** (16, 17). Most notably, the cytoplasmic complexes of all three systems appear as two inverted V’s in side view and as outer knobbed and inner contiguous rings in end-on view. The outer and inner rings are similar in cross-section (~225-300 nm) and height (~80 Å), suggestive of a common subunit composition. Supporting this proposal, structural analyses of mutant machines deleted of each of the T4SS ATPases, coupled with density tracing of a GFP moiety fused to a VirB4 homolog in the F system, strongly indicate that the inverted V structures of the F and Dot/Icm systems correspond to dimers of VirB4-like ATPases (16, 17). The cytoplasmic complexes of the F plasmid and Dot/Icm systems are therefore composed of VirB4 subunits arranged as a hexamer of dimers surrounding a central channel. The IMC of the *H. pylori* Cag T4SS is architecturally more complex than the IMCs of the F plasmid-encoded and Dot/Icm systems, yet VirB4-like Cagβ also arranges as a central hexamer of dimers at the entrance to the Cag translocation channel (18).

**Fig. 3.**
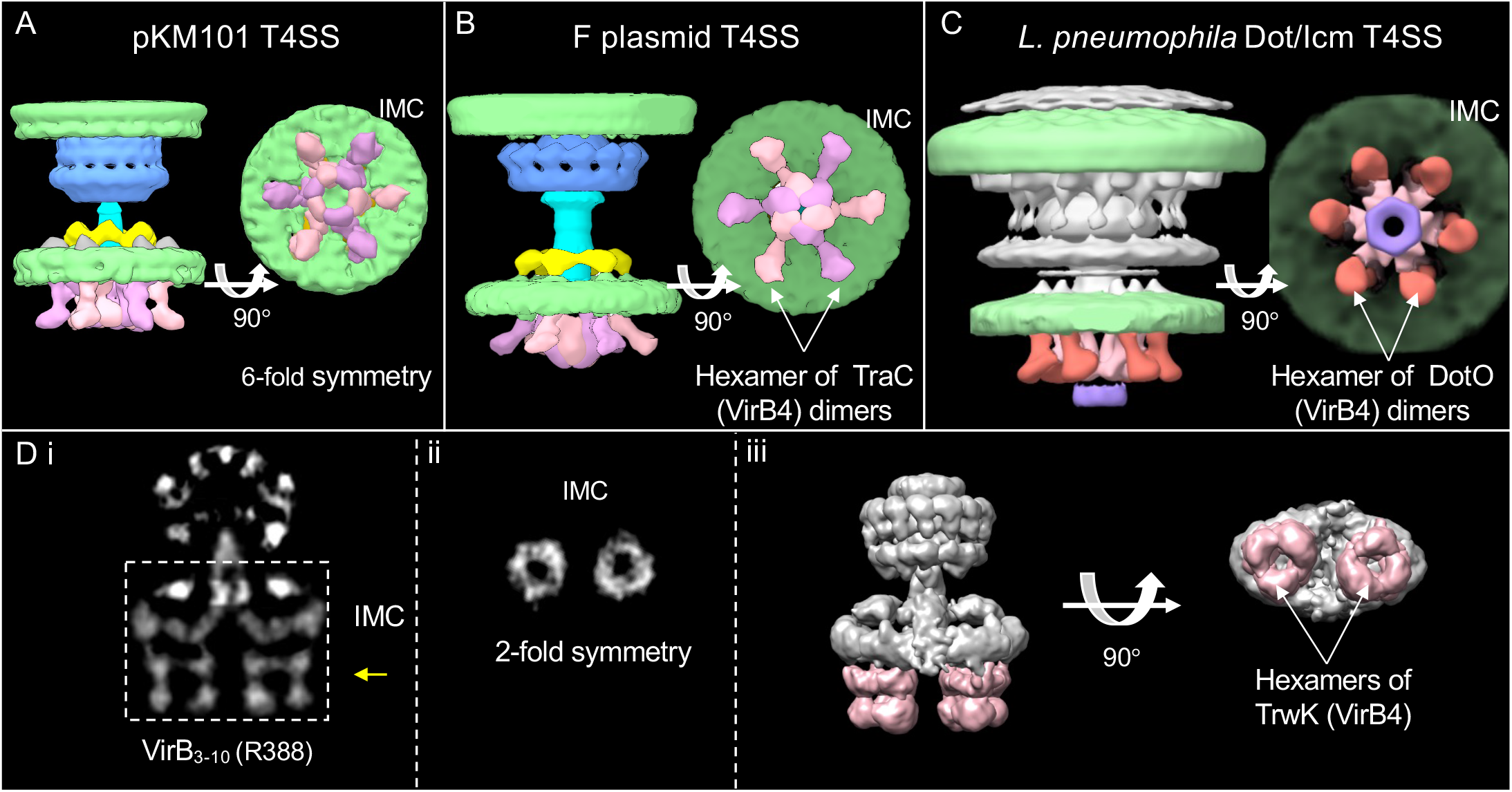
Comparison of the inner membrane complexes (IMCs) solved by CryoET and single particle analysis. (A, B, C) Comparison of the CryoET solved IMCs of type IV secretion systems encoded by pKM101 (T4SS_pKM101_), F plasmid (T4SS_pED208_) and *Legionella pneumophila* (T4SS_Dot/Icm_). 3D surface renderings showing 6-fold symmetric IMCs marked by the hexamer of dimer arrangement of VirB_4_ homologs. (D i) A central slice of the averaged structure of the purified VirB_3-10_ encoded by plasmid R388. (D ii) A cross-section view of the region in D i marked by a yellow arrow shows 2-fold symmetry exist in IMC of purified VirB_3-10_. (D iii) The surface rendering of the VirB_3-10_ substructure highlighting the IMC with side-by-side two hexamers of the TrwK/VirB4 ATPase (pink-shaded). F plasmid EMD: 9344 and 9347; Dot/Icm EMD: 7611 and 7612; VirB_3-10_ EMD:2567

The IMC_pKM101_ visualized *in situ* differs remarkably from the IMC of the isolated VirB_3-10_ structure **(Figs. 2B, C, 3A, 3Di-iii)** (6). In fact, in side view, the two complexes appear generally similar in the presence of side-by-side densities projecting into the cytoplasm. In end-on view, however, the collar of the *in situ* structure displays 6-fold symmetry, whereas the corresponding IM platform of the VirB_3-10_ structure is highly asymmetric. The inverted V’s visualized in the *in situ* structure form two concentric rings each with 6-fold symmetry, whereas the cytoplasmic complex of the *in vitro* structure is arranged as side-by-side barrel complexes (6). Each barrel comprises three tiers, and a central channel evident in the lower two tiers appears to be closed in the upper tier. By gold labeling, the barrels were shown to consist at least partly of the VirB4 homolog TrwK (6). A hexameric arrangement for the two barrels was inferred by earlier findings that TrwK assembles *in vitro* as a homohexamer, and results of stoichiometric analyses showing that the VirB_3-10_ complex is composed of 12 copies of TrwK_B4_ (6, 25).

### The pKM101 cytoplasmic complex is dominated by VirB4-like TraB

To test a prediction that VirB4-like TraB assembles as a hexamer of dimers at the entrance to the T4SS_pKM101_ channel, we imaged Δ*traB* mutant machines (100 machines from 254 tomographic reconstructions) **(Fig. S5).** *trans* expression of *traB_B4_* fully complements the Δ*traB* mutation, confirming that the mutation is nonpolar on downstream gene expression (8). All subvolume class averages of the Δ*traB* mutant machines showed the prominent OMCC but absence of discernible IM-associated densities (**Fig. S5**). Most notably, the Δ*traB* mutant machines lacked cytoplasmic densities dominated by the V structures (**Fig. 4B, S5**). The N-terminal domains (NTDs) of VirB4 ATPases associate tightly with the IM (26, 27), whereas C-terminal domains (CTDs) are highly conserved and adopt a RecA-like α/β structural fold implicated in ATP binding (28, 29). An atomic structure of the CTD of a VirB4 homolog fitted optimally within densities comprising the proximal halves of the V arms (**Fig. 4Eiii**). This is line with a proposal that the cytoplasmic complex of the *in situ* T4SS_pKM101_ is composed of TraB_B4_ configured as a hexamer of dimers, reminiscent of the architectures of the VirB4 homologs in the ‘expanded’ systems (16–18).

**Fig. 4.**
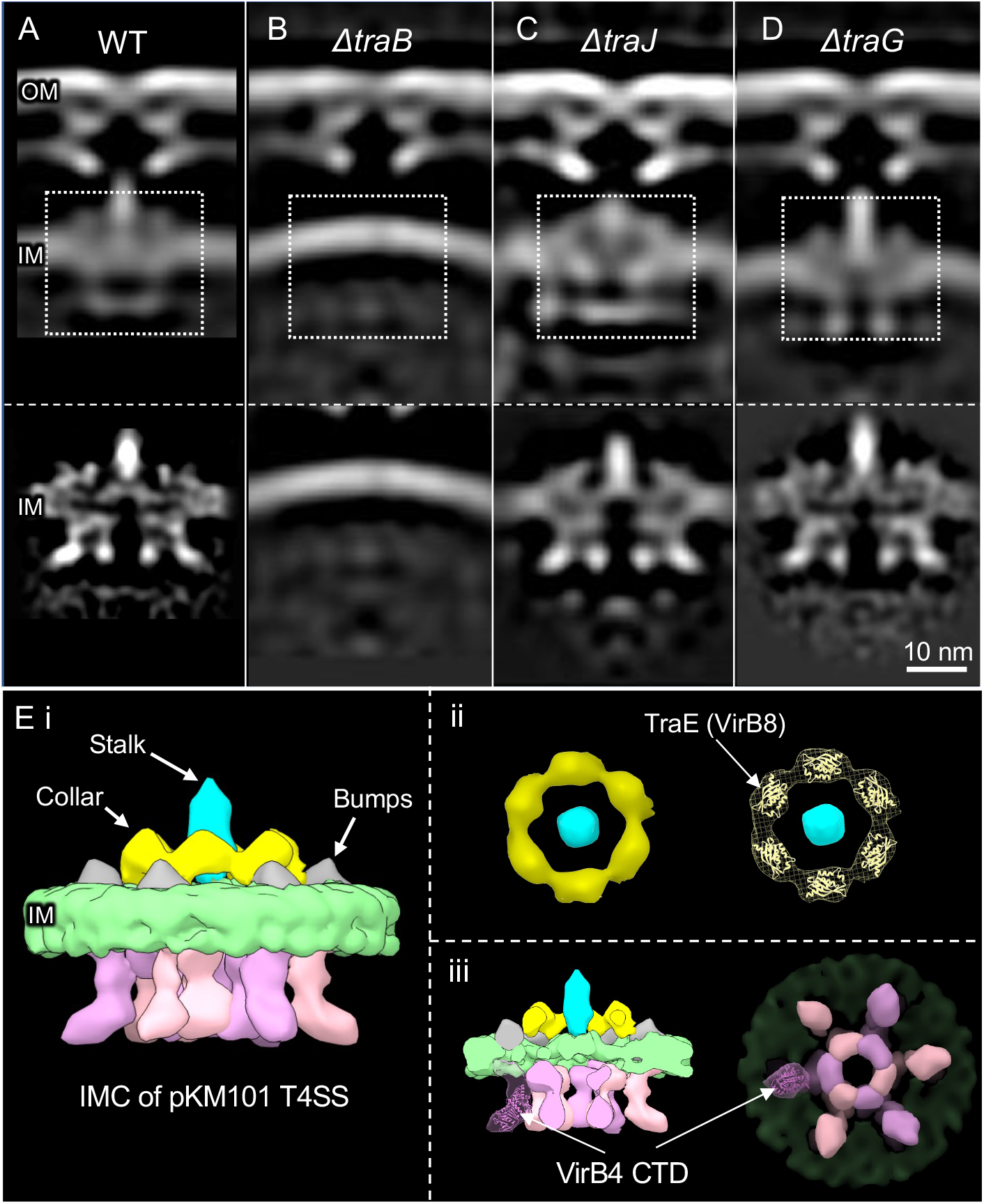
Architecture of the T4SS_pKM101_ machines from various ATPase mutants and comparison of the structure of the inner membrane complex (IMC) of pKM101 encoded type IV secretion system (T4SS_pKM101_) with solved structures. (A, B, C, D) The top row shows a central slice of the averaged structure of the T4SS seen in wildtype (WT), Δ*traB*, Δ*traJ* and Δ*traG* mutant cells. The second row shows the refined IMCs of the corresponding strains. (A) Native T4SS_pKM101_ machines (WT) for comparisons. (B) The Δ*traB* mutant machine lacks the periplasmic collar and cytoplasmic inverted “V” structures. (C, J) Mutant machines lacking TraJ and TraG align well with the WT structure. (E i) 3D surface renderings of the IMC of T4SS_pKM101_ is shown. (E ii) The collar exist as a hexameric ring in the periplasm just above the inner membrane (IM). Periplasmic domain of TraE_B8_ from pKM101 (PDB: 5I97) fits well in the lobe like structure of the collar. (E iii) Fitting of the C-terminal domain (CTD) of VirB4 ATPase from *Thermoanaerobacter pseudethanolicus* (PDB:4AG5) in one of the arms of hexamer of dimers of TraB.

Deletion of TraB_B4_ did not affect assembly of the OMCC **(Fig. 4B)**, but abolished other IMC densities including the periplasmic stalk and surrounding collar and ‘bumps’ (**Figs. 4B, S5**). VirB4 homologs associate peripherally with the IM (30) or at most possess small periplasmic domains (26), arguing against appreciable contributions of TraB_B4_ to the observed collar and stalk densities. Among the IMC subunits, the linker domains of TraF_B10_ and VirB5-like TraC_B5_ likely comprise the central stalk as suggested by compositional analyses of the VirB_3-10_ substructure (6). VirB8-like TraE also is a strong candidate for the central collar on the basis of recent nsEM studies showing that TraE_B8_ assembles as a homohexamer with dimensions matching those of the collar visualized *in situ* (31). An atomic structure of the periplasmic domain of TraE_B8_ from pKM101 (32) also fits well within each lobe of the collar (**Fig. 4Eii**). Finally, the polytopic VirB6 subunits typically carry large central periplasmic domains (33), which might form the ‘bumps’ or also contribute to the central collar or stalk domains. TraB_B4_ thus might need to be in place in order to recruit the IMC interaction partners to build out the IMC. In line with this proposal, among various characterized T4SSs, VirB4 subunits have been shown to form extensive networks of stabilizing interactions with VirB5, VirB8 and VirB10 homologs (8, 34–37).

### Deletions of VirD4-like TraJ or VirB11-like TraG do not detectably alter the *in situ* T4SS_pKM101_

Nearly all T4SSs require a VirD4-like ATPase to couple secretion substrates with the transfer channel. Designated as type IV coupling proteins (T4CPs) or substrate receptors, VirD4 subunits are members of the SpoIIIE/FtsK superfamily of motor translocases (38, 39). To determine if VirD4-like TraJ contributes to densities of the T4SS_pKM101_, we imaged Δ*traJ* mutant machines (183 machines from 430 tomographic reconstructions). As observed with the WT machines, subvolume averaging yielded classes of Δ*traJ* mutant machines exhibiting only the OMCC or both the OMCC and IMC densities (**Fig. S5**). A refined structure generated from the latter showed no distortions compared to WT machines, insofar as the OMCC, periplasmic collar and central stalk, and cytoplasmic V structures were clearly evident **(Fig. 4C).** The IMC thus clearly lacks detectable densities attributable to TraJ_D4_, which is reminiscent of findings for the F plasmid-encoded Tra and *L. pneumophila* Dot/Icm systems (16, 17). In the VirB_3-10_ structure, densities thought to correspond to one or two dimers of the VirD4 homolog were shown to embed between the VirB4 barrels (13). However, our *in situ* findings suggest the visualized densities likely represent transition-state structures or are artefacts resulting from machine purification. Evidence that VirD4-like subunits assemble and function as homohexamers (38, 40), and engage with T4SS channels only when activated by intracellular signals such as substrate engagement and ATP hydrolysis (41–45) further supports the notion that the T4CPs engage only transiently with the T4SS channel and therefore are difficult to detect by *in situ* CryoET.

Many T4SSs also require a third, VirB11-like ATPase for substrate transfer and pilus production (35, 46). VirB11 ATPases assemble as homohexamers that fractionate with the cytoplasm and IM, the latter presumably in association with the T4SS (16, 47–49). To determine if TraG_B11_ contributes to the visualized IMC_pKM101_, we imaged Δ*traG_B11_* mutant machines (257 machines from 537 tomograms). Of particular interest, in contrast to the WT and Δ*traJ_D4_* mutant machines in which IMC densities were detected in only ~50 % of the subvolume class averages, the fraction of Δ*traG* mutant machines with IMCs increased to 85 %. Furthermore, the visualized IMC densities were more clearly defined than in the class averages for the WT and Δ*traJ_D4_* mutant machines; in particular, the central stalks were considerably elongated (**Figs. 4, S5**). These findings suggest that TraG_B11_ might dynamically regulate assembly or conformational status of the central stalk and IMC as a function of ATP hydrolysis. This proposal is in line with previous work suggesting that VirB11 ATPases regulate the switch between two alternative structural modalities of T4SSs, the pilus biogenesis factory vs the active translocation channel (50). We also were unable to detect any density losses in the refined structure of the Δ*traG* mutant machine compared with the WT machine, further suggesting that TraG_B11_ regulates machine biogenesis through transient interactions with the T4SS_pKM101_ (**Fig. 4**). Consistent with this model, in the *L. pneumophila* Dot/Icm system, VirB11-like DotB shuttles dynamically between the cytoplasm and T4SS_Dot/Icm_ as a function of ATP hydrolysis (16). When engaged with the translocation channel, DotB_B11_ presents as a disc-shaped density attached to the inner hexameric ring of DotO_B4_. However, in order to visualize this density, it was necessary to deploy a DotB_B11_ mutant that bound but failed to hydrolyze ATP (16).

### Summary

CryoET has emerged as a critical complementary approach to single-particle CryoEM studies of bacterial secretion nanomachines (51). Although current resolutions achievable with CryoET are lower than CryoEM, structural definition of machines in their native contexts enables i) validation of architectural features observed *in vitro*, ii) assessments of machine structural variability within and between species, and iii) visualization of dynamic aspects of machine biogenesis and function (3, 51, 52). Here, we presented the first *in situ* structures of a ‘minimized’ T4SS, elaborated by the model conjugative plasmid pKM101. We showed that the T4SS_pKM101_ assembles *in vivo* as two large substructures, the OMCC and IMC, and that the former resembles structures of equivalent complexes solved *in vitro* (4, 7, 24). We further identified specific OM contacts and supplied evidence for OM remodeling during T4SS_pKM101_ biogenesis, and we visualized a central stalk similar to that detected in the isolated VirB_3-10_ complex (6). This latter finding establishes that the central stalk is an important *in vivo* structural element of ‘minimized’ T4SSs and was not a structural artefact of the isolated VirB3-10 complex resulting from detergent solubilization. Notably, the central stalk associated with the ‘minimized’ systems lacks a discernible central channel and thus is quite distinct from the corresponding structures connecting OMCCs and IMCs of the ‘expanded’ systems that are configured as larger cylinders with well-defined channels (**Fig. S4F**) (16–18).

Most importantly, we showed that the IMC of the *in situ* T4SS_pKM101_ nanomachine adopts an architecture completely different from that of the isolated VirB_3-10_ structure, especially with respect to the arrangement of the VirB4 ATPase (6). Rather than side-by-side hexameric barrels, VirB4-like TraB assembles as a central hexamer of dimers, an oligomeric state highly reminiscent of those of VirB4 homologs associated with the F-encoded Tra, *L. pneumophila* Dot/Icm and *H. pylori* Cag T4SSs (16–18). To reconcile these different results, we note that, in contrast to OMCCs, IMCs are characteristically highly unstable and difficult to purify in the presence of detergents, despite use of epitope-tagged subunits during purification such as VirB10-like subunits that stably associate with both subassemblies (4, 9–12, 24). In this context, it is reasonable to suggest that the VirB_3-10_ complex, and particularly its IMC, underwent structural distortions when stripped from the native membrane environment, which would render this portion of the reported structure largely an experimental artefact. Of course, it is feasible that the VirB_3-10_ complexes (6, 13) represent transition-state intermediates associated with machine assembly, but this seems unlikely since we were unable to detect such complexes in *in situ*.

Taken together with previous biochemical and structural data (21, 34, 35, 43, 53), our findings support an overarching model highlighting contributions of the VirB4 ATPases to biogenesis of T4SSs regardless of their classification as ‘minimized’ and ‘expanded’ (**Fig. S6**). Assembly initiates by formation of the intrinsically-stable OMCC (**stage I**), which for the T4SS_pKM101_ system can be visualized *in situ* in the absence of associated densities. The OMCC then recruits VirB4 (**stage II**), most likely through previously-identified interactions between the ATPase and the cell-envelope-spanning VirB10 subunit (35). Once positioned, VirB4 recruits other components of the IMC, including subunits such as VirB5, VirB6, and VirB8 that likely contribute to the visualized periplasmic collar and flanking ‘bumps’, and central stalk structures. Upon receipt of an unknown signal, VirB4 then recruits the spatially-dynamic VirB11 ATPase (**stage III**), which induces structural changes in the stalk and IMC of postulated importance for the pilus to channel transition and channel opening (21, 50). Finally, the T4SS channel is activated by a combination of extracellular (target cell contact) and intracellular (substrate docking, T4CP engagement with the T4SS channel, ATP energy utilization) signals, resulting ultimately in the passage of substrates through the VirB4 hexamer and into the envelope-spanning channel (**stage IV**). While there has been remarkable recent progress in defining the architectures of T4SSs, a central question remains how the VirD4 substrate receptor associates with the T4SS and conveys substrates into the channel. Given that this is likely a transient interaction, further deployment of *in situ* CryoET approaches aimed at capturing structural snapshots of T4SSs - including those in the act of translocating substrates - will provide valuable new insights into structural transitions accompanying machine activation.

## Materials and Methods

### Strains and growth conditions

Bacterial strains, plasmids, and oligonucleotides used in this study are listed in Table S1. *E. coli* strains were grown at 37°C in Luria Bertani (LB) agar or broth supplemented with appropriate antibiotics (kanamycin, 100 μg ml^−1^; spectinomycin, 100 μg ml^−1^; gentamycin, 10 μg ml^−1^). Minicells from *E. coli* strain UU2834 were used for all of the CryoET studies.

### Conjugation assays

*E. coli* MG1655 strains carrying pKM101 or mutant variants were used as donors to transfer the plasmids into UU2834 recipients. Strains containing the pKM101 mutants also harbored a complementing plasmid. Overnight cultures of donor and recipient cells grown in presence of the appropriate antibiotics at 37 °C were diluted 1:1,000 in fresh LB media, and incubated with shaking for 1.5 h. When needed, cells were induced with arabinose (0.2 % final concentration) and incubated with shaking for another 1.5 h. Equal volumes (50 μl) of donor and recipient cell cultures were mixed and incubated for 3 h at 37 °C. Mating mixtures were serially diluted and plated onto LB agar containing antibiotics selective for transconjugants (Tc’s). Plasmid-carrying UU2834 strains were verified for the presence or absence of *tra* genes of interest by PCR, and used for CryoET. For matings to assess minicell donor capacity, minicells were spotted onto a nitrocellulose filter disc alone or with MC4100*rif^r^* recipient cells, and the mating mixes were incubated at 37°C for 1 h. The disc was suspended in LB, serially diluted and plated on LB agar plates containing appropriate antibiotics selective for Tc’s and donors (to confirm absence of viable donor cells).

### Isolation of minicells

*E. coli* minicells were enriched essentially as described previously (54). *E. coli* UU2834 harboring pKM101 or variants were grown overnight at 37°C in LB in the presence of spectinomycin, and then subcultured (1:100) in fresh LB devoid of antibiotics at 37°C to an OD_600_=0.5. Anucleate minicells were selectively enriched by centrifugation at 1,000 *x g* for 3 min, the supernatant was centrifuged again at 1,000 *x g* for 3 min, and minicells were passed through a 0.45 μm filter. Minicells were then used in mating assays. To minimize possible breakage of the pKM101-encoded pilus and to concentrate minicells for CryoET analyses, UU2834 strains were grown overnight on LB agar plates at 37 °C. Cells were gently scraped from the plate surface with an “L” shaped rod and resuspended in phosphate-buffered saline (PBS). The cell suspension was centrifuged twice at 1,000 *x g* for 3 min to remove intact cells and the supernatant was then centrifuged at 20,000 *x g* for 20 min and the minicell pellet was resuspended in PBS for preparation of grids for CryoET.

### Preparation of frozen-hydrated specimens

Minicells resuspended in PBS were mixed with 10 nm diameter colloidal gold particles and deposited onto freshly glow-discharged, holey carbon grids for 1 min. After blotting the grids with filter paper, they were rapidly frozen in liquid ethane by using a gravity-driven plunger apparatus (55, 56).

### Cryo-ET data collection and 3D reconstructions

Frozen-hydrated specimens were imaged and data were processed using our previously established protocols (12, 28, 41). Briefly, specimens were subjected to imaging at −170°C using a Polara G2 electron microscope (FEI Company) equipped with a field emission gun and a direct detection device (Gatan K2 Summit). The microscope was operated at 300 kV at a magnification of 15K, resulting in an effective pixel size of 2.5 Å at the specimen level as previously described (17). Tomographic package SerialEM (57) was used to collect low-dose, single-axis tilt series with dose fractionation mode with defocus at about 6 μm and a cumulative dose of ~60 e^−^/Å^2^ distributed over 35 stacks. Each stack contains ~8 images. Each tilt series was collected at angles from −51° to 51° with 3° fixed increment. We used Tomoauto (55) to expedite data processing, which included drift correction of dose-fractionated data using Motioncorr (58) and assembly of corrected sums into tilt series, automatic fiducial seed model generation, alignment and contrast transfer function correction of tilt series by IMOD (59), and reconstruction of tilt series into tomograms by Tomo3D (60). Each tomographic reconstruction was 3,710 by 3,838 by 2,400 pixels and ~130 Gb in size.

### Subtomogram averaging and correspondence analysis

Tomographic package I3 (61) was used for subtomogram analysis as described previously (62). A total of 837 T4SS_pKM101_ machines (400 × 400 × 400 voxels) were visually identified and then extracted from 1781 cryo-tomographic reconstructions. Two of the three Euler angles of each T4SS_pKM101_ machine were estimated based on the orientation of each particle in the cell envelope. To accelerate image analysis, 4 × 4 × 4 binned subtomograms (100 × 100 × 100 voxels) were used for initial alignment. The alignment proceeded iteratively, with each iteration consisting of three parts in which references and classification masks are generated, subtomograms are aligned and classified, and, finally, class averages are aligned to each other. At the initial iterations, classification mask was applied to include the whole machine and non-T4SS particles were sorted out and removed. For analysis of the IMC, a mask was applied to the IMC only, thus the T4SS particles that did not show IMC density were sorted out and the data set showing IMC was used to further refine the IMC. Classification focusing on the OMC displayed 14-fold symmetry; therefore, 14-fold symmetry was imposed in the following processing to assist the initial alignment process. Furthermore, classification focusing on the IMC showed a 6-fold symmetry feature, and in the following processing 6-fold symmetry was imposed to assist in sub-tomograms alignment. After multiple cycles of alignment and classification for 4×4×4 binned sub-tomograms, 2×2×2 binned sub-tomograms was used for refinement. Fourier shell correlation (FSC) between the two independent reconstructions was used to estimate the resolution of the averaged structures (**see Fig. S2**).

### 3D visualization

IMOD was used to visualize the maps and generate 3D surface rendering of *E. coli* minicells. UCSF Chimera (63) (http://www.rbvi.ucsf.edu/chimera) was used to visualize subtomogram averages in 3D and for molecular modeling. The video clips for the supplemental videos were made by using UCSF Chimera and further edited with iMovie.

### Data availability

Density maps and coordinate data of the T4SS_pKM101_ machines determined by cryo-electron tomography have been deposited in the Electron Microscopy Data Bank (EMDB) as EMD-24100 and EMD-24098. The authors declare that all other data supporting the findings of this study are available within the paper and its supplementary information files.

## LEGEND FOR SUPPLEMENTARY MATERIALS

**Table S1.** List of strains, plasmids, and oligonucleotides used in these studies.

**Fig. S1.** Genes and encoded functions of “minimized” type IV secretion systems (T4SSs) in Gram-negative species.

**Fig. S2.** Workflow for *in situ* CryoET.

**Fig. S3.** Detection of pKM101 pilus docked on *E. coli* outer membrane.

**Fig. S4.** Comparisons of the outer membrane core complexes (OMCCs) and stalks/cylinders of ‘minimized’ and ‘expanded’ T4SSs.

**Fig. S5.** Heterogeneity of the pKM101 T4SS machines revealed by class sub-volume averaging.

**Fig. S6.** A unified model for T4SS assembly and routing of secretion substrates through central channels.

**Movie S1.** 3-D visualization of a tomographic reconstruction and the T4SS_pKM101_ in *E. coli* minicells.

**Movie S2.** 3-D visualization of the T4SS_pKM101_ showing architectural features of the OMCC and IMC and its comparison with purified VirB_3-10_ complex from R388.

## ACKNOWLEDGEMENTS

B.H. was supported by McGovern Medical School start-up funds, the Welch Foundation (AU-1953-20180324), NSF grant #1902392, and NIH 1R35GM138301. P.J.C. was supported by NIH 1R35GM131892. B. H. and P.J.C were supported by NIH R21AI142378. We thank members of the Christie and Hu labs for helpful discussions.

